# Presence and Stability of SARS-CoV-2 on Environmental Currency and Money Cards

**DOI:** 10.1101/2021.08.23.457328

**Authors:** Colleen Newey, Abigail T. Olausson, Alyssa Applegate, Ann Aubrey Reid, Richard A. Robison, Julianne H. Grose

## Abstract

The highly contagious nature of SARS-CoV-2 has led to several studies on the transmission of the virus. A little studied potential fomite of great concern in the community is currency, which has been shown to harbor microbial pathogens in several studies. Since the onset of the COVID-19 pandemic, many businesses in the United States have limited the use of banknotes in favor of credit cards. However, SARS-CoV-2 has shown greater stability on plastic in several studies. Herein, the stability of SARS-CoV-2 at room temperature on banknotes, money cards and coins was investigated. In vitro studies with live virus suggested SARS-CoV-2 was highly unstable on banknotes, showing an initial rapid reduction in viable virus and no viral detection by 24 hours. In contrast, SARS-CoV-2 was far more stable on money cards with live virus detected after 48 hours. Environmental swabbing of currency and money cards on and near the campus of Brigham Young University supported these results, with no detection of SARS-CoV-2 RNA on banknotes, and a low level on money cards. No viable virus was detected on either. These preliminary results suggest that the use of money cards over banknotes in order to slow the spread of this virus may be ill-advised. These findings should be investigated further through larger environmental studies involving more locations.

## Introduction

Coronavirus disease 2019 (COVID-19) was first reported in late December 2019 in Wuhan, Hubei Province, China[1–3] The disease is caused by severe acute respiratory syndrome coronavirus 2 (SARS-CoV-2), a beta coronavirus, and presents as an acute respiratory infection.[1, 2] Clinical presentation of COVID-19 commonly includes fever, cough, chest tightness, dyspnea, pneumonia, and fatigue, with loss of taste and smell also being reported.[4] According to molecular analyses, the beginning of the 2019 epizootic breakout of SARS-CoV-2 occurred around November 25th, 2019. On March 11, the World Health Organization made the assessment that the SARS-CoV-2 outbreak could be characterized as a pandemic and as of August 17, 2021 there have been over 4.3 million global deaths from COVID-19, with over 620,000 deaths in the United States alone (https://coronavirus.jhu.edu/map.html).

Coronaviruses are large, enveloped, positive-strand RNA viruses that belong to the *Coronaviridae* family. They primarily infect a wide range of animals and are most ecologically diverse among bats, with viruses from the *Alphacoronanvirus* and *Betacoronavirus* genera known to infect humans. The SARS-CoV-2 virus likely originated from the SARS-like virus in the *Rhinolophus* bat family[3] and falls within the *Sarbecovirus* subgenus.[5, 6] Though human coronavirus infections are typically mild, SARS-CoV, MERS, and presently SARS-CoV-2, all *Betacoronaviruses*, are notably severe exceptions to this rule.[7–9] Although SARS-CoV-2 is the least deadly, it is more contagious than SARS-CoV or MERS. Coronaviruses contain four major structural proteins: the spike surface glycoprotein (S), small envelope protein (E), matrix protein (M), and nucleocapsid protein (N). S proteins from different coronaviruses determine host specificity and allow entry into the host cells via different cell surface receptors. Like SARS-CoV, structural and molecular analyses have implicated ACE2 as a receptor target for SARS-CoV-2.[6, 10, 11]

To curb future outbreaks, both public health and molecular perspectives are essential to understand *SARS-CoV-2* transmission patterns. Although primarily transmissible from person-to-person through respiratory droplets, evidence shows the virus is also transmissible through direct contact, feces, or other body fluids.[12–15] *SARS-CoV-2* has been shown to be transmissible by asymptomatic carriers even during its incubation period.[16, 17] This may be due to the higher upper respiratory viral load observed in COVID-19 cases, as infection in the upper respiratory tract is more easily spread by coughing and sneezing.[18] Fomites, inanimate objects capable of absorbing, harboring and transmitting infectious microorganisms, have also been implicated as possible sources of transmission.[19] Several pathogens, including both gram-positive (*Clostridioides difficile*, *Enterococcus spp*., *Staphylococcus aureus* and *Streptococcus pyogenes*) and gram-negative (*Acinetobacter spp*., *Escherichia coli*, *Klebsiella spp*. *Pseudomonas aeruginosa*, *Serratia marcescens* and *Shigella spp.*) bacteria as well as some viruses (*Astrovirus, HAV, Poliovirus and Rotavirus*), have been reported to survive for months on surfaces while others are much less stable (reviewed by Kramer, Schwebke and Kampf).[20] Van Doremalen et al. released a study in March 2020 on the stability of *SARS-CoV-2* on various surfaces.[4] Five environmental conditions were tested including aerosol, plastic, stainless steel, copper and cardboard. Aerosolized virus was viable for at least 3 hours with only a 16% reduction in titer. Viable virus was detected on stainless steel and plastic for up to 72 hours, however, large reductions in titer were observed (from 10^3.7^ per mL media down to 10^0.6^ per mL media).[4] In contrast, no viable virus was detected on cardboard after 24 hours or on copper after 4 hours.[4]

In environmental, real world studies, a variety of substances and surfaces have been tested for the presence of *SARS-CoV-2* RNA, including aerosols[5, 21–28] and environmental surfaces[5, 22, 23, 25, 26, 28–30] in hospitals throughout the world, surfaces in a quarantine hotel room[31] as well as on chopsticks[32] in China, and a study of ferryboat, nursing homes, and COVID-19 isolation wards in Greece[33], with positive detection of viral RNA on a variety of surfaces in all studies with the exception of three of the aerosol studies[23, 25, 27] (recently reviewed by Meyerowitz[34]). Lack of detection in some aerosol studies has been attributed to differences in sample collection which may result in inactivation of the virus.[21] Live virus was assessed in three of these studies and detected in only two.[21, 22, 28] In the few studies of laboratory surfaces, *SARS-CoV-2* was either undetectable[35] or detected at low levels[36, 37].

A little studied potential fomite of great concern in the community is currency[38, 39] , which resulted in a halt of currency use in many businesses during the COVID-19 pandemic. Previously, investigators have outlined the suspected role of paper money and coins as bacterial disease vectors, including the detection of *S. aureus* on banknotes recovered from hospitals, and *S. aureus*, *Salmonella* and *E. coli* detected on banknotes recovered from food outlets.[40] Lower denominations contained more bacteria, consistent with higher circulation rates.[40] In a recent study by Harbourt et al., the stability of *SARS-CoV-2* was modeled in the laboratory on four common surfaces including swine skin, uncirculated United States of America $1 and $20 Federal Reserve notes, and scrub cloth, which served as a common clothing example.[41] At room temperature (22°C), they reported stability of the live virus after 4 but not 8 hours on clothing, after 8 hours but not 24 on $1 U.S.A. banknote, while the virus remained quantifiable up to 24 hours on the $20 U.S.A. banknote or skin. The virus was much more stable at 4°C. However, the stability of *SARS-CoV-2* on credit card or coin has not been reported, and there has been a lack of environmental testing outside of the laboratory.[42] In order to better understand the risk of paper money versus credit cards as fomites for *SARS-CoV-2*, the stability of the virus on U.S.A $1 banknotes, coin and money cards in the laboratory is studied herein. In addition, the frequency of detection of *SARS-CoV-2* on environmental paper money and money cards was assessed in the Brigham Young University (BYU), Provo, Utah area.

## Methods

### Regulations

For environmental samples, all environmental positive samples collected in 50% DMEM were disposed of in Biohazard bins after the addition of an equal volume of 70% ethanol. All assayed money cards were immediately disinfected after use with 70% ethanol. All work involving live SARS-CoV-2 was performed in the biosafety level-3 facility at BYU, Provo, UT. All work was conducted under proper safety handling procedures approved by the BYU Institutional Biosafety Committee (IBC-2020-0068).

### Cell Culture

Viral cultivation and plaque assays were performed using African green monkey kidney (VERO 76, C1008) cells obtained from American Type Culture Collection (ATCC). VERO cells were maintained in T-75 flasks containing Dulbecco’s Modified Eagle’s Medium (DMEM; Corning, 10-017-CV) supplemented with 10% Fetal Bovine Serum (FBS; Corning, 35-010-CV) at 37°C, and 5% CO_2_. For 24-well plate preparation, cells were seeded at 200,000 cells per well in DMEM with 10% FBS and used for assays 18-24 hours later.

### Viral Strain

SARS-CoV-2 (2019-nCoV/USA-WA1/2020) was obtained from the ATCC. Viral titer was obtained by plaque assay in VERO cells. Briefly, stock virus was diluted 1:2 and from 10^−1^ through 10^−10^ by 10-fold serial dilutions in DMEM. VERO cells in 24-well plates were then inoculated with 200ul of a virus dilution. Plates were placed at 37°C and 5% CO_2_ for 1 hour and manually rocked every 10 minutes. A 1ml-overlay of 1:1 mixture of 2X Minimal Essential Medium (MEM; Hyclone, SH30008.04) with 8% FBS and 1.5% low melting agarose (ThermoFisher Scientific, R0801) was placed in the wells. Agarose plugs were allowed to solidify at room temperature for 30 minutes. Plates were then incubated for 72 hours at 37°C and 5%CO_2_. Post incubation, wells were fixed with 10% formaldehyde for 1 hour. Agar plugs and formaldehyde were then removed and wells were washed twice with distilled water. 200ul of crystal violet stain (0.4%) were then placed in each well for 3 minutes at room temperature, followed by rinsing with distilled water twice and air drying. Plaques were counted and titers were performed in duplicate.

### Purification of SARS-CoV-2 RNA (positive control for LAMP assays)

SARS-CoV-2 infected VERO cells were lysed by adding 0.3-0.4 mL of TRIzol Reagent (Invitrogen, ThermoFisher Scientific) per 1 × 10^5^-10^7^ cells directly in the culture dish and pipetting up and down several times to homogenize. Samples were allowed to incubate for 5 minutes, and then centrifuged for 5 minutes at 12,000 ×g at 4–10°C. The clear supernatant was transferred to a new tube and stored at −80 °C.

### Environmental U.S. Currency Experiments

Due to the reduction in usage of cash in and around campus, environmental samples of cash were primarily obtained from the Brigham Young University vault, a daily collection of currency and coin from university-based stores, restaurants, dormitories and vending machines. In addition, fresh samples were obtained from local restaurants and assayed within one hour. The entire surface of cash and/or coin was swabbed around the edges of the bill and then back and forth across the surface for ~10 seconds with a sterile cotton swab moistened with sterile 50% DMEM (1X with 4.5 g/L glucose, L-glutamine & sodium pyruvate from Corning, Cat# 10-013-CV diluted into sterile molecular grade water). The swab was immersed with agitation in 500uL 50% DMEM. The samples were immediately assayed for viral signatures within one hour after collection using a loop-mediated isothermal amplification (LAMP) assay. The WarmStart LAMP Kit (Cat# E1700S, New England BioLabs) was utilized using SARS-CoV-2 specific primers[43] or SARS-CoV-2 Rapid Colorimetric LAMP Assay Kit (New England Biolabs), which became available in assays after September 15, 2020. Primers, described by El-Tholoth et al., were diluted to a final concentration of: 16 uM FIP, 16 uM BIP, 2uM F3, 2uM B3, 4uM LOOP F, 4 uM LOOP B[44]. Samples were incubated at 65°C for 60 minutes and then cooled on the bench top for ten minutes before recording color. To verify positives, samples were also analyzed by DNA gel electrophoresis using a 1% agarose gel and ethidium bromide. All positive samples from the initial assays were immediately assayed a second time and only those that appeared positive by either the colorimetric indicator or by gel electrophoresis with characteristic bands were counted as true positives.

### Lab Inoculation of U.S Currency Experiments

Viable SARS-CoV-2 virus (20ul of stock at 1.5×10^7^ pfu/mL) was inoculated onto designated U.S currency; $1 U.S.A. banknotes, credit card, quarter, and penny. Currency was sampled at the following four post-inoculation time points: 0, 4, 24, and 48 hours. Negative controls were inoculated with 20ul of sterile DMEM. The initial time point post treatment began at media drying point on U.S.A. banknotes (~30 minutes) in an operating biosafety cabinet at 22°C. At each time, cotton swabs were dipped in 500ul DMEM contained in 1.5 ml microcentrifuge tubes and used to swab area where virus was inoculated. Swabs were immediately placed back into the collection microcentrifuge tube of 500ul media and mixed thoroughly to extract virus. Samples were then serially diluted; 50ul sample into 450ul DMEM from 10^−1^ to 10^−4^ VERO cells in 24-well plates were inoculated with 200ul of each dilution; Undiluted, 10^−1^, 10^−2^, 10^−3^, 10^−4^, and DMEM negative control. Plates were placed at 37°C and 5% CO_2_ for 1 hour and manually rocked every 10 minutes. A 1ml-overlay of a 1:1 mixture of Minimal Essential Medium (MEM; Hyclone, SH30008.04) and 1.5% low melting agarose (ThermoFisher Scientific, R0801) maintained at 37°C was placed in wells. Agarose plugs were allowed to solidify at room temperature for 30 minutes. Plates were then incubated for 72 hours at 37°C and 5% CO_2_. Post incubation, wells were fixed with 10% formaldehyde for 1 hour. Agar plugs and formaldehyde were then removed, and wells were washed twice with distilled water. 200ul of crystal violet stain (0.4%) were placed in each well for 3 minutes at room temperature, followed by rinsing with distilled water twice and air drying. Plaque counts were then determined. All tests were performed in duplicate. Previously circulated $1 U.S.A. banknotes, credit cards and coins were utilized and were sterilized by U.V. light prior to use in this study.

### Local Coronavirus Case Counts

Local coronavirus case counts were obtained from https://coronavirus.utah.gov/case-counts/.

### Statistical Analysis

Graphs were produced using GraphPad Prism software version 8.0.0 (GraphPad Software, San Diego California) and ANOVA with post hoc Tukey HSD was performed using JMP pro version 15 (JMP®, Version *15.* SAS Institute Inc., Cary, NC, 1989-2021). Half-life was estimated from the plot of the log_2_ of the titer, using JMP Pro to fit a line and deriving the half-life from the slope according to the equation (−log_2_ 2/slope). Time points after a time point approximating a zero titer were dropped from the linear fit.

## Results

### Laboratory testing of SARS-CoV-2 reveals instability on banknotes but improved stability on money cards

Viable SARS-CoV-2 was spotted on $1 banknotes, quarters, pennies and credit cards and then extracted by swabbing with a MEM-moistened sterile swab at time points 30 minutes (allowing time for drying) as well as 4, 24, and 48 hours post inoculation. Extracted samples were immediately assayed for viable SARS-CoV-2 via plaque counts in VERO cells (Fig. 1). SARS-CoV-2 was difficult to extract from $1 banknotes, even immediately (30 minutes) after inoculation, with a 10^5^-fold or 99.9993% reduction in titer (2.5×10^6^ to 1.75×10^1^ pfu/mL, p = 8.66 x10^−4^). Further significant reductions in viable virus occurred at the 24 and 48 hour time points, where no live virus was detected (p = 0.012 and 0.039, respectively when compared with the 30 minute time point). In contrast, money cards displayed only a 10-fold or 90% reduction in titer at 30 minutes (2.5×10^6^ to 2.5×10^5^ pfu/mL, p = 1.51 ×10^−3^) which may in part be due to a difference in the ability of the viral suspension to soak into or bind plastic money cards versus paper banknotes. Further significant reductions in titer occurred at 4 hours (99.6% reduction compared with 30 minute values, p = 1.7 x10^−7^), however live virus was still detectable at 24 and 48 hours post inoculation, displaying a 99.96% and 99.97% reduction, respectively, when compared with time 30 minutes (p = 1.2 x10^−7^ and p = 7.9 x10^-7^, respectively). Quarters and pennies were similar to the money card, with a stronger initial reduction in viral titer (99.4% and 99.6%, respectively, at time 30 minutes, p = 8.9 x10^−4^ and 3.6 x10^−3^) and virus detectable at the 24 and 48 hour time points. Thus, viable SARS-CoV-2 appeared to be most stable on plastic money cards, with banknotes providing the least stability of all four surfaces tested in this study.

**Fig. 1.**
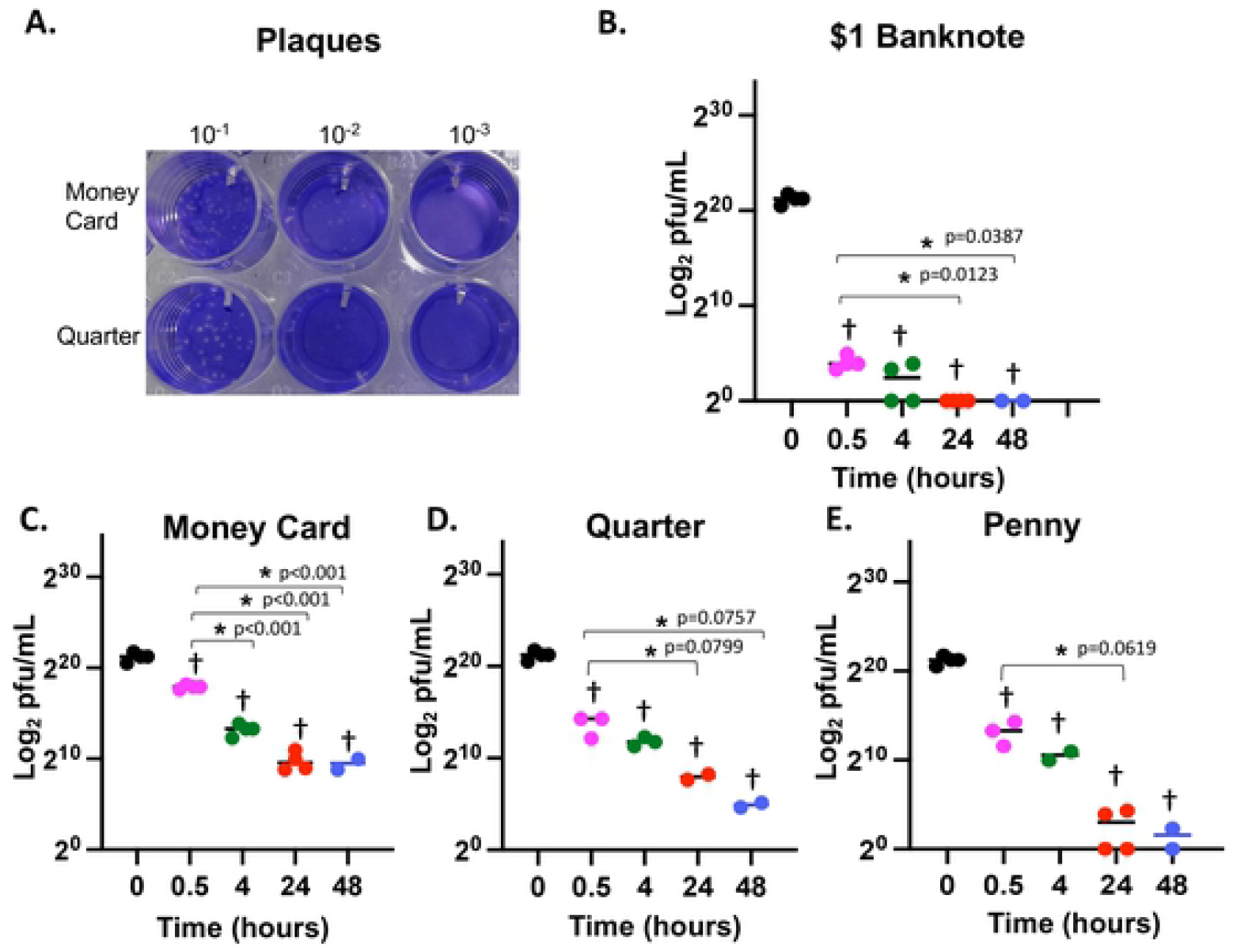
Stability of live SARS-CoV-2 virus on $1 U.S.A. banknotes, money cards, quarters, and pennies. A) Representative plaques of a SARS-CoV-2 on VERO cells from this study at the four hour time point. B) Recovery of virus after inoculation of $1 U.S.A. banknotes after 0.5, 4, 24 and 48 hours. C) Recovery of virus after inoculation of money cards after 0.5, 4, 24 and 48 hours. D) Recovery of virus after inoculation of quarters after 0.5, 4, 24 and 48 hours. E) Recovery of virus after inoculation of pennies after 0.5, 4, 24 and 48 hours. Log_2_ of the virus concentration was represented on the y-axis of all charts to better display differences in numbers. At zero titer, log_2_ is undefined and is therefore shown at 2^0^. Time 0.5 was used as an initial time point after inoculation to allow time for drying on each surface. ANOVA followed by post hoc Tukey HSD using JMP Pro (JMP®, Version *15.* SAS Institute Inc., Cary, NC, 1989-2021) was used to detect differences between time points, and all significant differences (p< 0.10) between the time 0.5 hours and other time points as indicated by “*”. For every surface, the starting point (just prior to inoculation) was significantly different (p<0.05) from all other time points, indicated by “†“.

All samples displayed a rapid, initial drop in viable virus (PFU/mL), followed by a different rate of decay. This may be due to surface drying times. Therefore, modeling of the half-life of SARS-CoV-2 on these surfaces was conducted with and without inclusion of the starting titer to obtain two-step decay rates, initial rates (from time 0 to time 0.5 hours) of decay versus later rates (from time 0.5 hours onward, Table 1). The initial apparent half-life of SARS-CoV-2 was shortest on banknotes (~1.7 minute half-life), quarters (~ 4.0 minutes) and pennies (~ 4.0 minutes) compared to the money card (~ 9 minutes) (Table 1 and Fig. 1). Second step decay rates were again highly variable, with quarters displaying slower decay than money cards or pennies, however money cards decay appeared to level out at 24 hours while no or extremely low levels of SARS-CoV-2 were detected on pennies after 24 hours. The $1 U.S.A. banknote data for second step decay calculations is unreliable due to variability the detection of extremely low levels of virus at the four-hour time point (0 versus 10 or 15 PFU/mL), suggesting an almost complete loss of titer at the 30 minute time point (see Fig. 1).

**Table 1.** Apparent two-step half-life of SARS-CoV-2 on various forms of currency.

| Half Life* | \$1 U.S.A. Banknote | Money Card* | Quarter | Penny |
| --- | --- | --- | --- | --- |
| Second-stage half-life | 4.3 hour<br>(0.530) | 3.3 hour<br>(0.769) | 5.6 hour<br>(0.933) | 2.5 hour<br>(0.66) |
| Initial Half-life | 0.029 hour<br>(0.997) | 0.15 hour<br>(0.958) | 0.066 hour<br>(0.963) | 0.068 hour<br>(0.983) |

\*Two-step half-life was estimated from the plot of the $\log_2$ of the titer (Plaque Forming Units, PFU), using JMP Pro to fit a line and deriving the half-life from the slope according to the equation $(-\log_2 2/\text{slope})$ . $R^2$ is given in parenthesis. Due to nonlinearity, time points after an approximately zero titer were dropped from the linear fit. For all samples, the initial half-life was calculated from the starting titer and the 30 minute time point (0.5 hours). For the second-stage half-life, the 30 minute time point was used as the starting time point. Time points after 24 hours were dropped for the \$1 U.S.A. Banknote, money card, and penny due to their low titers (approximately zero). For the quarter, all time points (excluding time zero) were utilized in the second stage calculation. The \$1 U.S.A. banknote second-stage half-life is unreliable due to almost complete loss of virus by time 30 minutes.

### SARS-CoV-2 RNA was detected on environmental money card samples but not on paper money

Due to the apparent instability of SARS-CoV-2 on U.S. banknotes and the improved stability on money cards as assayed in the laboratory, a large-scale screening was performed to detect SARS-CoV-2 RNA on environmental samples. Two hundred and seventy-nine money cards, including credit cards and Brigham Young University (BYU) ID money cards, were swabbed and assayed via SARS-CoV-2 LAMP assay (Fig. 2, Table 2). A total of 17 money cards tested positive in duplicate LAMP assays. However, a positive by LAMP assay indicates the presence of SARS-CoV-2 RNA and not live virus. To determine if there was live virus present, a total of 14 samples from the 2/19/2021 assays (including all 6 samples that were positive) were interrogated in the BSL-3 laboratory for live virus using a standard plague assay. No plaques were seen for any of the samples. Dilutions of a viable suspension of SARS-CoV-2 with a titer of 7.75×10^6^ were used as positive controls, suggesting that any positive LAMP tests were likely due to SARS-CoV-2 RNA only.

**Fig. 2.**
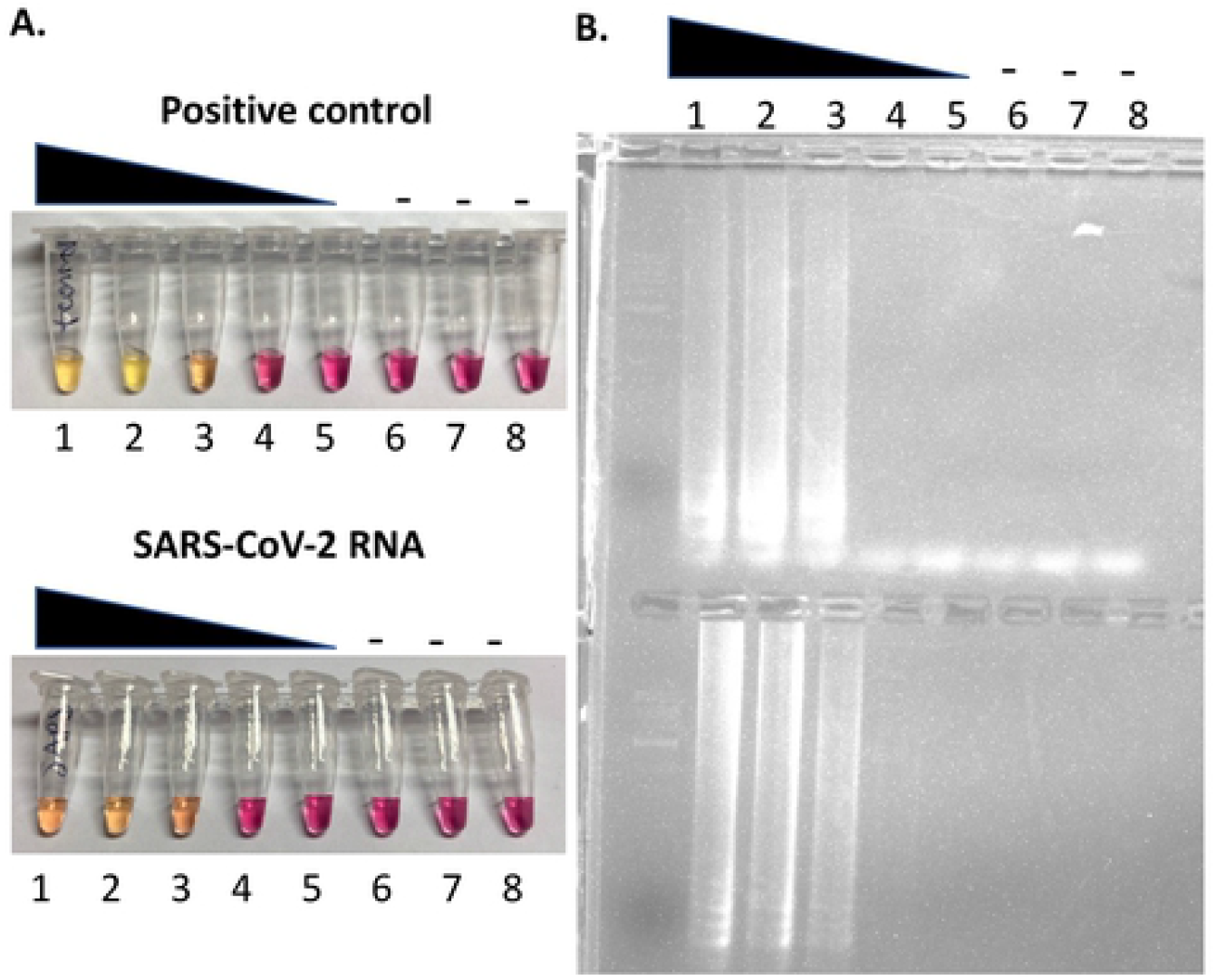
LAMP assay-based detection of SARS-CoV-2. A) Representative colorimetric LAMP SARS-CoV-2 assay using the SARS-CoV-2 Rapid Colorimetric LAMP Assay Kit (New England Biolabs) and NEB positive control or purified SARS-CoV-2 RNA. B) Associated gel electrophoresis of samples from (A). NEB positive control and SARS-CoV-2 RNA were used straight (sample number 1), and then serially diluted 1:10, four times (sample numbers 2-5). Samples 6-8 are buffer only negative controls.

**Table 2.** LAMP assays for detection of SARS-CoV-2 RNA on money cards on Brigham Young University (BYU) campus.

| **Date collected** | **COVID cases per 100,000 (daily)\*** | **# assayed** | **# LAMP positive (%)\*\*** | **# Plaque assay positive (%)** |
| --- | --- | --- | --- | --- |
| 11/17/2020 | 122.3 (79.4 ) | 110 | 8 (7.2%) | - |
| 12/11/2020 | 92.1 (99.8) | 90 | 3 ( 3.3%) | - |
| 2/19/2021 | 29.4 (22.8) | 79 | 6 (7.6%) | 0*** |
| **Totals:** |  | 279 | 17 (2.6%) |  |

\*Active, laboratory confirmed, 7-day average COVID cases from Utah County as declared from the local health department (daily rate in parenthesis).
\*\*Positive samples are positives from an initial assay that were verified in a secondary assay.
LAMP assays were conducted using the SARS-CoV-2 Rapid Colorimetric LAMP Assay Kit (New England Biolabs).
\*\*\*A total of 14 samples, including all 6 positive samples and 8 negatives from the 2/19/2021 testing were assayed in the BSL-3 laboratory for live virus using a dilution series starting at 1:4.
No plaques were seen for any of the samples. A viable suspension of SARS-CoV-2 with a titer of $7.75 \times 10^6$ pfu/ml was used as a positive control. “-“ is not determined.

Four hundred and twenty-nine U.S.A. banknotes from the BYU campus vault (or nearby restaurants) were also tested for the presence of SARS-CoV-2 RNA. No sampled banknotes tested positive, consistent with the observed instability observed on banknotes (Table 3). Only one hundred and five coins were tested due to the lack of coin circulation at the time of testing and none were positive.

**Table 3.** LAMP assays for detection of SARS-CoV-2 RNA on environmental U.S.A. banknotes and coins.

| Date(s)<br>Collected | Date<br>Sampled | Source | COVID cases per<br>100,000(daily)* | #<br>Banknotes<br>Sampled<br>(# coins) | # LAMP<br>Positive** |
| --- | --- | --- | --- | --- | --- |
| 6/11/2020-6/12/2020 | 6/12/2020 | BYU vault | 8 (9) | 73 (27) | 0 |
| 6/16/2020-6/17/2020 | 6/17/2020 | BYU vault | 9.3 (14) | 222<br>(78) | 0 |
| 10/14/2020 | 10/14/2020 | Restaurants<br>near BYU | 51.7 (64.8) | 134 | 0 |
| **Totals:** |  |  |  | 429(105) | 0 |

\*Active, laboratory confirmed, 7-day average COVID cases from Utah County as declared from the local health department (daily rate in parenthesis).
\*\*Positive samples are positives from an initial assay that were verified in a secondary assay. LAMP assays were by WarmStart Colorimetric LAMP (New England Biolabs) or SARS-CoV-2 Rapid Colorimetric LAMP Assay Kit (New England Biolabs) which became available for the 10/14/2020 testing.

## Discussion

In the United States, paper currency is a blend of ~75% cotton and 25% linen with ample surface area, which is routinely passed between individuals suggesting that it may function as a fomite in the spread of many bacterial and viral diseases. When the coronavirus pandemic was announced, a common reaction from businesses was to stop all use of U.S.A. banknotes in order to hypothetically reduce the spread of this virus. Neither the World Health Organization (WHO) nor the Centers for Disease Control (CDC) in the United States has banned the use of paper money and coin, but they have emphasized that people should wash their hands after handling money as a precaution. Many stores are still either limiting or completely denying the use of cash. Our results suggest that SARS-CoV-2 is far more stable on plastic money cards, with only a 99.6% reduction of the virus after four hours, compared with a 99.9993% reduction of the virus on $1 U.S.A. banknotes immediately after inoculation of U.V.-sterilized, circulated banknotes (Fig. 1). These results agree with those of Harburt et al., who reported no detection of SARS-CoV-2 at 24 hours post inoculation of uncirculated $1 U.S.A. banknotes at room temperature.*[41]* In addition, we were unable to detect some SARS-CoV-2 RNA on U.S.A. banknotes in circulation (Table 3), consistent with the instability of SARS-CoV-2 on banknotes in vitro. Also consistent with in vitro assays, we were able to detect SARS-CoV-2 RNA on environmental money cards at a low level (Fig. 1 and Table 2). However, no live virus was detected on any of the environmental money card samples. These results suggest that the use of money cards over banknotes is not advisable, although these preliminary results should be confirmed with larger studies over several locations.

Our results are consistent with reports that SARS-CoV-2 is far more stable on plastic than paper, and support the recommendation that paper sheets and bags be used instead of plastic to reduce the spread of COVID-19.[4, 41, 45–48] In a large-scale study of banknotes from various countries, polymer-based banknotes (such as those used in New Zealand and Australia) have been shown to display lower bacterial counts than cotton-based banknotes, and in regions such as Mexico, where both types of banknotes are utilized, fewer counts were observed on polymer-based notes. Corpet has proposed this is due to the dryness of paper and other porous solids.[46] This hypothesis would be consistent with the results we observed where the inoculum on paper money displayed a huge initial (time 30 minute) decline in titer, which may be due to an initial drying on or binding to the paper. In contrast, money cards did not show this drastic initial decrease, but instead showed a gradual decrease to 24 hours, followed by low level stability which may be due to increased time necessary for drying on the surface. However, the hypothesis that the decrease in stability is due to dryness of the paper and porous solids is unproven, and appears to contradict the experimental studies showing higher reduction in coronaviruses numbers at high humidity (80% versus 20%).[46] Thus, instability on paper money may reflect SARS-CoV-2 binding properties to the surface composition rather than humidity/dryness. Clearly further study is necessary to understand the binding and stability properties of SARS-CoV-2.

## Acknowledgements

We thank the Department of Microbiology and Molecular Biology and the College of Life Sciences at Brigham Young University for their support. In addition, we thank the Brigham Young University Treasury Services, including Steven Morley and Tammy Miner, as well as local businesses and numerous students at Brigham Young University for providing samples for this study.

